# Elevated GLUT4 Levels in Human Skeletal Muscle Microtissues is Accompanied by Functional Insulin Dependence

**DOI:** 10.1101/2025.04.24.650438

**Authors:** Yekaterina Tiper, Jiaru Ni, Roman Krawetz, Penney M Gilbert

## Abstract

Insulin resistance in skeletal muscle is a hallmark of type 2 diabetes mellitus (T2D). While two-dimensional myotube cultures offer a controlled environment for studying T2D-related metabolic dysfunction, insulin-dependent glucose transporter type 4 (GLUT4) levels are limited and insulin-independent glucose transporter type 1 (GLUT1) expression dominates; reducing physiological relevance. Three-dimensional skeletal muscle microtissue cultures offer a promising alternative, and unlike 2D myotubes, are amenable to repeated contractile stimulation. However, microtissue GLUT1 and GLUT4 glucose transporter profiles remain under-characterized, particularly under physiological glucose and insulin conditions, which is evaluated herein. We report that GLUT1 levels trended ∼3.0-fold lower in microtissues compared with myotubes in 2D culture, although not statistically significant (*p* = 0.072), while GLUT4 levels were ∼12-fold higher (*p* < 0.0001), leading to a ∼60-fold increase in the GLUT4:GLUT1 ratio (*p* = 0.023). Notably, the microtissue GLUT4:GLUT1 profile approached, but did not match that of native human muscle. Microtissues required supraphysiological insulin conditions for the development of maximal contractility, while physiological glucose levels were sufficient. Insulin withdrawal restored insulin responsiveness but impaired microtissue contractile strength (*p* < 0.0001) and fatigue resistance (*p* = 0.015). Our findings indicate that the glucose transporter profile of microtissues offers improved physiological relevance. However, their reliance on insulin to maintain contractile function limits their suitability for modeling T2D. The implementation of a robust, insulin-free differentiation protocol would facilitate the development of a microtissue-based T2D model which can be applied to study contraction-mediated increases in insulin sensitivity as a therapeutic approach.

## 1. INTRODUCTION

Type 2 diabetes mellitus (T2D) is a metabolic disorder characterized by impaired insulin secretion from pancreatic β-cells and systemic insulin resistance. Skeletal muscle, as the primary site of glucose disposal, is one of the earliest tissues affected in T2D and plays a critical role in driving insulin resistance (Merz & Thurmond, 2020). The tissue maintains glucose homeostasis through both insulin-dependent and insulin-independent mechanisms. Under basal conditions, glucose transporter type 1 (GLUT1) primarily mediates glucose uptake, ensuring a steady supply of glucose to muscle cells (Ciaraldi et al., 2005). Postprandial glucose uptake is predominantly driven by glucose transporter type 4 (GLUT4), which translocates to the cell membrane in response to insulin stimulation (Lund et al., 1997). Independently of insulin, GLUT4 is also recruited to the membrane during contraction-mediated glucose uptake in response to exercise, providing an alternative pathway for glucose regulation (Douen et al., 1990; Kennedy et al., 1999). Defective insulin-stimulated GLUT4 translocation is a major contributor to insulin resistance in skeletal muscle, leading to reduced glucose uptake and hyperglycemia (Klip et al., 2019).

As the incidence of T2D continues to rise globally, there is an urgent need to elucidate the molecular mechanisms underlying skeletal muscle dysfunction in diabetes and develop more effective therapeutic strategies. *In vitro* skeletal muscle models have become a key tool in this pursuit, providing a controlled environment to investigate T2D-related metabolic dysfunction. Among these, induced T2D models enable the study of specific pathological features by exposing cultured myotubes to elevated levels of glucose, insulin, lipids, and/or inflammatory factors (Hirabara et al., 2010; Steinberg et al., 2006; Turner et al., 2020). Alternatively, subject-derived models capture cell-autonomous diabetic phenotypes by utilizing primary or induced pluripotent stem cell (iPSC)-derived myoblasts from individuals with T2D (Batista et al., 2020; Carcamo-Orive et al., 2020; Kase et al., 2015).

While two-dimensional (2D) myotube-based T2D models have provided valuable insights into both the intrinsic and extrinsic contributors to insulin resistance and metabolic dysfunction, their physiological relevance remains limited. 2D cultures fail to adequately support the contractile properties of skeletal muscle, as myotubes lack proper alignment and detach from the substrate upon reaching contractile maturity, impeding their differentiation potential. Moreover, they exhibit metabolic deficiencies, including an altered glucose transporter profile characterized by low GLUT4 expression and high GLUT1 levels (Sarabia et al., 1992). Since GLUT4 is the primary mediator of insulin-stimulated glucose uptake in skeletal muscle, its reduced expression in 2D cultures compromises insulin responsiveness, and limits the utility of 2D cultures in modeling T2D.

To overcome the limitations of traditional 2D myotube cultures, three-dimensional (3D) skeletal muscle models have emerged as a more physiologically relevant alternative (Martin et al., 2013; Vandenburgh et al., 2008). These models are generated by seeding myoblasts within an extracellular matrix and differentiating them under uniaxial tension between two fixed points, thereby mimicking the mechanical environment of native muscle. Unlike 2D cultures, engineered muscle forms aligned, contractile myotubes, providing a platform to investigate how contraction and contraction-induced cytokines (i.e., myokines) enhance insulin sensitivity (Benrick et al., 2012; Chen et al., 2020; Holten et al., 2004). Notably, engineered skeletal muscle has demonstrated greater insulin responsiveness than 2D myotubes derived from the same cell source (Kondash et al., 2020). However, engineered muscle glucose transporter profiles relative to native muscle remain uncharacterized, highlighting a gap in our understanding of its metabolic properties.

Despite these advances, a model of T2D in engineered skeletal muscle has yet to be established. One major challenge lies in the reliance on differentiation protocols that often incorporate supraphysiological levels of insulin, glucose, and/or amino acids to promote myoblast fusion and contractile myotube formation (Hofemeier et al., 2021; Madden et al., 2015; Mills et al., 2019). Differentiation under hyperglycemic and hyperinsulinemic conditions induces metabolic adaptations that render models unresponsive to additional glucose or insulin. This limitation prevents the study of extrinsic mechanisms driving glucose and insulin dysregulation. Therefore, optimizing differentiation conditions to more closely mimic the physiological metabolic environment, while preserving the contractile function of engineered muscle, is essential for establishing a translationally relevant *in vitro* model of T2D.

Previously, we reported MyoTACTIC, a 96-well platform for the bulk generation of human skeletal muscle microtissues (hMMTs), enabling contractile force assessment via micropost deflection (Afshar et al., 2020). In this study, we investigated the glucose transporter and insulin receptor content of hMMTs and 2D myotubes derived from the same primary cell line, relative to native human muscle. We also assessed whether hMMTs could be differentiated in physiologically relevant culture media without compromising contractile function.

## 2. MATERIALS AND METHODS

### 2.1 Human primary myoblast culture and 2D differentiation

A human primary myoblast line was obtained from Cook MyoSite Inc. (SK-1111-P01358-19F; Pittsburgh, PA, USA), which derived using a proprietary method from the vastus lateralis muscle of a 19-year-old Caucasian female. The donor had a body mass index of 24 kg/m^2^ and no known medical conditions. Myoblasts were passaged as previously described (Gulati et al., 2024; Tiper et al., 2025) In brief, they were maintained on rat tail collagen I (Cat. #A1048301, Gibco; Waltham, MA, USA) coated tissue culture plates. Cells were propagated in growth medium composed of Ham’s F-10 nutrient mix (Cat. #318-050-CL, Wisent Bioproducts; Saint-Jean Baptiste, QC, Canada) supplemented with 20 % fetal bovine serum (FBS; Cat. #12483020, Gibco), 5 ng/mL basic fibroblast growth factor (bFGF; Cat. #11343625, ImmunoTools; Friesoythe, Germany), and 1 % Penicillin-Streptomycin (P/S; Cat. #15140122, Gibco). The medium was refreshed every other day. Upon reaching ∼70 % confluency, myoblasts were detached from the culture surface using Trypsin-EDTA (0.25 %; Cat. #25200072, Gibco) and replated at a lower density. At passage 8, cells were used for differentiation into 2D myotubes and for the fabrication of hMMTs. The population was ∼95 % CD56^+^ at this passage, as determined by fluorescence-activated cell sorting, indicating a high degree of myogenic purity.

For 2D differentiation, myoblasts were plated on rat tail collagen I coated 48-well and 6-well plates at ∼70,000 cells/cm^2^. Cells were maintained in growth medium for 1-2 days before transitioning to differentiation medium. Differentiation was induced using high-glucose (4.5 g/L) Dulbecco’s Modified Eagle’s Medium (DMEM; Cat. #11995065, Gibco) or Minimum Essential Medium (MEM; Cat. #11095080, Gibco) supplemented with 2 % horse serum (HS; Cat. #16050114, Gibco), and 1 % P/S, with or without 1.72 µM insulin (10 µg/mL; Cat. #I6634, Sigma-Aldrich; Burlington, MA, USA). Half the medium was refreshed every other day. Endpoint analyses were performed after 4-5 days of differentiation. Myotubes differentiated in 48-well plates were used for immunostaining and imaging, while those differentiated in 6-well plates were used for western blotting.

### 2.2 MyoTACTIC culture platform fabrication

The polydimethylsiloxane (PDMS; Sylgard^TM^ 184 silicone elastomer kit, Dow Corning; Midland, MI, USA) MyoTACTIC 96-well culture platform was fabricated following previously reported methods (Afshar et al., 2020; Lad et al., 2021). In brief, all features of the plate were cast in a single step from a reusable polyurethane negative mold. After casting, the PDMS plate was sectioned into portions of 4 ± 2 wells, then sonicated in isopropanol for 20-30 minutes, rinsed with dH_2_O, dried in a curing oven at 60-65 °C, and autoclaved in an instrument sterilization bag. To create a non-adhesive culture surface conducive to tissue remodelling, PDMS portions were incubated overnight at 4 °C with a 5 % Pluronic® F-127 solution (diluted in dH_2_O; Cat. #P2443, Sigma-Aldrich). The solution was aspirated immediately prior to hMMT seeding.

### 2.3 hMMT seeding and differentiation

hMMTs were seeded in the MyoTACTIC culture platform as previously described (Afshar et al., 2020; Lad et al., 2021). In brief, human primary myoblasts were suspended in a hydrogel mixture of 4 mg/mL fibrinogen (40 % v/v; Cat. #F8630, Sigma-Aldrich) and Geltrex^TM^ (20 % v/v; Cat. #A1413202, Gibco) in high-glucose DMEM. The cell suspension ranged from 1.00 x 10^7^ to 1.67 x 10^7^ cells/mL, corresponding to 1.50 x 10^5^ to 2.50 x 10^5^ cells/tissue. Thrombin (0.37 units/mg of fibrinogen; Cat. #T6884, Sigma-Aldrich) was added to initiate polymerization, and 15 µL of the cell-hydrogel suspension was quickly dispensed into each PDMS culture well. After 5 min of polymerization at 37 °C, after-seeding growth medium, consisting of Ham’s F-10 nutrient mix with 20 % FBS, 1.5 mg/mL 6-aminocaproic acid (ACA; 3 % v/v; Cat. #A2504, Sigma-Aldrich), and 1% P/S, was added. hMMTs were maintained in this medium for two days before transitioning to differentiation medium. Differentiation was induced by culture in high-glucose DMEM, MEM, or low-glucose (1.0 g/L) DMEM (Cat. #11885084, Gibco) supplemented with 2 % HS, 2 mg/mL ACA (4 % v/v), and 1 % P/S, either without or with up to 1.72 µM insulin, as specified in the text. Endpoint analyses were conducted after 14 days of differentiation. A summary of all media and solutions used for human primary myoblast expansion, 2D myotube differentiation, and hMMT production and culture can be found in **Supplemental Table 1**.

### 2.4 hMMT electrical field stimulation and evaluation of hMMT contractile force

To assess hMMT contractile function at endpoint, electrical field stimulation (EFS) was applied in the MyoTACTIC culture platform as previously described (Afshar et al., 2020; Lad et al., 2021). In brief, hMMTs were stimulated between two needle electrodes (25G x 5/8”; Cat. #305122, BDTM; Franklin Lakes, NJ, USA) inserted into the well at either end of microtissue. Micropost deflection during EFS was recorded using an iPhone SE (Apple®; Cupertino, CA, USA) mounted on the eyepiece of an Olympus IX83 inverted microscope (Tokyo, Japan) with the 10X objective. Videos were analyzed using a semi-automated Python script available on GitHub (Gilbert Lab, 2024), which tracks a selected region of the deflecting micropost across frames and calculates total post displacement during contractions. Displacement values in pixels were converted to microns using a pixel-to-distance conversion factor, then to contractile force in Newtons using a distance-to-force conversion factor.

EFS was performed at the optimized stimulation parameters eliciting peak force (Tiper et al., 2025). To quantity dynamic oscillation of force (DOF), 6-7 contractions were recorded at 1 Hz, 5 V, and 60 ms. DOF, a metric we previously introduced for evaluating the contractile force of engineered skeletal muscle produced from human primary myoblasts, is defined as the difference between nadir and peak force following contraction stabilization at 1 Hz. It was calculated by averaging values from the 3^rd^, 4^th,^ and 5^th^ contractions in the stimulus train (Tiper et al., 2025).After a 30-second rest period, tetanus force and fatigue resistance were assessed. Five contractions were recorded at 7 Hz, 5 V, and 10 ms. Tetanus force was defined as the maximum absolute force achieved during the 1^st^ tetanic. Fatigue resistance was calculated as the force generated during the 6^th^ tetanic contraction, expressed as a percentage of that achieved in the 1^st^ contraction.

### 2.5 Immunohistochemistry and image acquisition

To visualize myotubes at endpoint, the media from 2D cultures and 3D hMMTs was aspirated, and samples were washed with Dulbecco’s Phosphate-Buffered Saline (D-PBS). Fixation was performed in 4 % paraformaldehyde (PFA; Cat. #50-980-494, Electron Microscopy Sciences; Hatfield, PA, USA) in D-PBS for 15 min at room temperature (RT), followed by three washes in D-PBS. For 2D myotubes, cultures were permeabilized for 15 min in 0.1 % Triton X-100 (Cat. #TRX777, BioShop; Burlington, ON, CA) and 1 % bovine serum albumin (BSA; Cat. #ALB005, BioShop) in D-PBS, then blocked with 10 % goat serum (GS; Cat. #16210072, Gibco) in D-PBS for 1 hour at RT. Cultures were incubated overnight at 4°C with mouse anti-sarcomeric α-actinin (SAA) primary antibody (1:800; Cat. #A7811, Sigma-Aldrich) in PBS with 1 % GS. The next day, cultures were washed three times with 0.025 % Tween-20 (Cat. #TWN510, BioShop) in PBS and incubated for 1 hour at RT in a secondary antibody mix consisting of Alexa Fluor^TM^ 488 goat anti-mouse secondary antibody (1:500; Cat. #A11001, Invitrogen; Waltham, MA, USA), and Hoechst 33342 nuclear counterstain (1:1000; Cat. #H3570, Invitrogen) in PBS with 1 % GS. Three final washes with 0.025 % Tween-20 in D-PBS were performed before imaging. hMMTs were permeabilized, blocked, and stained as previously described (Afshar et al., 2020; Ebrahimi et al., 2021), using the same primary and secondary antibody mixture as for 2D myotubes. In brief, hMMTs were blocked and permeabilized in 10 % GS with 0.3 % Triton X-100 in D-PBS for 1 hour at RT, then incubated overnight at 4 °C with the primary antibody diluted in the blocking/permeabilization solution. The next day, hMMTs were washed three times with D-PBS and incubated for 1 hour at RT in the secondary antibody mixture diluted in the blocking/permeabilization solution. Three final washes with D-PBS were performed prior to imaging.

Confocal z-stack images of immunostained samples were acquired using an Olympus IX83 inverted microscope at a step size of 5 µm and processed with Olympus Fluoview-10 software. A 20x and 40x objective was used to image 2D myotubes and hMMTs, respectively. For 2D myotubes, 5 randomly selected regions per well were imaged. For hMMTs, 3-4 non-overlapping images were captured along the length of each microtissue.

### 2.6 Nuclear fusion index and SAA coverage analysis for 2D myotubes

To assess myoblast fusion in 2D cultures, nuclear fusion index and SAA coverage were quantified by analyzing images of the SAA-immunostained and Hoechst-counterstained myotubes. For nuclear fusion index analysis, confocal z-stacks of the SAA and Hoechst channels were projected at maximum intensity and merged using Fiji (ImageJ) (Schindelin et al., 2012). The multi-point tool was used to count nuclei within myotubes, defined as SAA-positive cells containing more than two nuclei, as well as the total number of nuclei in the field of view. Fusion index was calculated as the percentage of nuclei within myotubes relative to the total nuclei count. For SAA coverage analysis, the confocal z-stack of the SAA channel was projected at maximum intensity and thresholded using the Huang method (Huang & Wangt, 1995) to distinguish foreground from background. SAA coverage was determined as the percentage of foreground pixels relative to the total number of pixels in the image. Nuclear fusion index and SAA coverage percentages were averaged across 5 images per well.

### 2.7 Western blotting

The levels of GLUT4, GLUT1, and insulin receptor β (INSRβ) in 2D and 3D cultures were assessed via western blotting. To compare protein levels in these cultures with those in native muscle, a human skeletal muscle biopsy was included as a reference in the western blot analysis. The sample was derived from gracilis skeletal muscle tissue collected within 4 hours postmortem from a 64-year-old male donor with no history of arthritis, joint injury, or surgery, no prescription anti-inflammatory medications, and no comorbidities. Informed consent was obtained from next of kin. The collection and use of the human skeletal muscle tissue was reviewed and approved by the University of Calgary Research Ethics Board (REB15-0005). The University of Toronto Office of Research Ethics also reviewed the approved this study and assigned administrative approval (Protocol# 34888). All procedures in this study were conducted in accordance with the guidelines and regulations of both Research Ethics Boards.

Protein samples were prepared by pooling 1–2 wells of myotubes differentiated in a 6-well plate for 2D cultures and 3-5 hMMTs for 3D cultures. Samples were transferred to 1.5 mL microtubes and homogenized on ice in lysis buffer containing 25 mM Tris (pH 7.2; Cat. #TRS001, BioShop), 0.5 % Triton X-100, and a protease and phosphatase inhibitor cocktail (1:100; Cat. # 78440, Thermo Scientific; Waltham, MA, USA). Homogenization was performed using a handheld electric motor (Cat.#FS7495400000, Fisherbrand; Waltham, MA, USA) with disposable pellet pestles (Cat. #FS7495211590, Fisherbrand). Lysates were centrifuged at 1500 x g for 10 min at 4°C, and the supernatants, containing the soluble sarcoplasmic proteins, were transferred to fresh 1.5 mL microtubes (Roberts et al., 2020). Protein concentrations of the sarcoplasmic isolates were determined using a bicinchoninic acid (BCA) assay (Cat. #23227, Thermo Scientific), and samples were stored at -80 °C until use.

Samples were prepared for sodium dodecyl sulfate-polyacrylamide gel electrophoresis (SDS-PAGE) by adding NuPAGE^TM^ LDS sample buffer (25 % v/v; Cat. #NP0007, Invitrogen) and NuPAGE^TM^ sample reducing agent (10 % v/v; Cat. #NP0009, Invitrogen) to 15 ug of sarcoplasmic protein isolate. Samples were heated at 70 °C for 10 min in a water bath, followed by incubation with 100 mM iodoacetamide (10 % v/v; Cat. #IOD500, BioShop) for at least 30 min at RT covered from light. After brief centrifugation, samples were loaded onto NuPAGE^TM^ 4-12% Bis-Tris mini protein gels (Cat. #NP0321BOX, Invitrogen). Protein separation was performed at 80 V for ∼2.5-3 hours in NuPAGE^TM^ MOPS SDS running buffer (Cat. #NP000102; Invitrogen) supplemented with NuPAGE^TM^ antioxidant (Cat. #NP0005, Invitrogen). Proteins were transferred from the polyacrylamide gel to a nitrocellulose membrane at 60 V for 1 hour using a wet transfer system. The transfer buffer consisted of 25 mM Tris, 192 mM glycine (Cat. #GLN001, BioShop), and 20 % (v/v) methanol. Membranes were then stained with Ponceau S solution (Cat. #PON002, BioShop) to visualize total protein content before being blocked for 1 hour in Tris-buffered saline with 0.1% (v/v) Tween-20 (TBST) containing 5 % (w/v) skimmed milk (Cat. #SKI400, BioShop). Blocked membranes were incubated overnight at 4 °C with primary antibodies directed against GLUT4 (1:1000; Cat #PA1-1065, Invitrogen), GLUT1 (1:100; Cat. #sc-377228, Santa Cruz Biotechnology; Dallas, TX, USA), or INSRß (1:200; Cat. #sc-57342, Santa Cruz Biotechnology) diluted in TBST with 5 % skimmed milk. The next day, membranes were washed three times in TBST (≥5 min per wash) and incubated for 1 hour at RT with horseradish peroxidase (HRP)-conjugated secondary antibodies, anti-rabbit (Cat. #7074, Cell Signaling Technology; Danvers, MA, USA) or anti-mouse (Cat. #7076, Cell Signalling Technology), both diluted 1:5000 in TBST with 1 % skimmed milk. Following three additional TBST washes, membranes were incubated with enhanced chemiluminescent substrate (Cat. #A38555, Thermo Scientific) for HRP conjugate detection. Images were acquired using a ChemiDoc imaging system (BioRad; Hercules, CA, USA), and band intensities were quantified using ImageLab software (BioRad). All intensities were normalized to total protein content. A list of antibodies and the counterstain used in this study, along with the host species, working dilutions, and manufacturer information can be found in **Supplemental Table 2**.

### 2.8 Quantification of 2-deoxyglucose uptake

For glucose uptake experiments, hMMTs were differentiated in low-glucose DMEM supplemented with 0.86 µM insulin for 10 days. Following differentiation, hMMTs were either maintained in high-glucose DMEM supplemented with 0.86 µM insulin (as control) or cultured in low-glucose DMEM without added insulin to induce insulin re-sensitization. On day 14, insulin-stimulated glucose uptake was assessed in control and re-sensitized hMMTs using the glucose analog 2-deoxyglucose (2DG), as previously described for engineered skeletal muscle (Kondash et al., 2020). This method relies on the phosphorylation of 2DG to 2-deoxyglucose-6-phosphate (DG6P) upon cellular uptake, which is quantified using an enzymatic assay that produces resorufin fluorophore in proportion to DG6P levels (Yamamoto et al., 2015).

In brief, on the day of the assay, hMMTs were first washed with D-PBS, then serum- and insulin-starved for 4 hours. After an additional D-PBS wash, hMMTs were incubated for 30 min in uptake buffer consisting of glucose-free DMEM (Cat. #11966025, Gibco) supplemented with 0.1% BSA and 0.2 mM sodium pyruvate (Cat. #11360070, Gibco). Insulin-stimulated and basal glucose uptake were assessed by incubating hMMTs for 50 minutes in uptake buffer with or without 200 nM insulin. During the final 20 minutes of incubation, 1 mM 2DG (Cat. #D6134, Sigma-Aldrich) was added to the media.

To terminate the reaction, hMMTs were washed three times with D-PBS and placed immediately on ice. Each hMMT was carefully transferred from the MyoTACTIC platform into a 96-well microplate using forceps, with one tissue placed per well. For sample lysis and NADP elimination, 50 µL of 0.1 M NaOH was added to each well, and the plate was heated at 65-75 °C for ∼45 min. The samples were then neutralized with 50 µL of 0.1 M HCl, followed by the addition of 50 µL of 200 mM triethanolamine (TEA; Cat. #T1502, Sigma-Aldrich) buffer, and mixed on a shaker. For DG6P detection, 25 µL of each sample was transferred to a white opaque microplate (Cat. #15042, Thermo Scientific) and incubated with 200 µL of assay solution **(Supplemental Table 3)** at 37 °C for 60 min. Fluorescence was measured using a Tecan microplate reader (λex = 530 nm, λem = 590 nm; Infinite 200 Pro; Männedorf, Switzerland).

### 2.9 Statistical analysis

The number of wells or hMMTs serving as technical replicates (n) and the number of biological replicate experiments (N) are specified in the figure legends. Data are reported as the mean ± standard deviation (SD), and error bars represent SD in bar graphs. Statistical significance was assessed using an unpaired, two-tailed *t*-test for two-group comparisons, and a one-way ANOVA for multiple-group comparisons, followed by Dunnett’s or Tukey’s post hoc test as appropriate. A significance threshold of *p* ≤ 0.05 was applied to all tests. Details regarding technical replicates, pooled tissue counts, and statistical tests for each experiment are provided in **Supplemental Table 4**. All graphs were generated, and statistical analyses were performed using GraphPad Prism 10.0 (GraphPad Software; Boston, MA, USA).

## 3. Results

### 3.1 hMMTs exhibit an elevated GLUT4:GLUT1 ratio compared to 2D myotubes

To investigate whether glucose transporter and insulin receptor content is modulated by nutrient and insulin conditions during myotube differentiation, we quantified GLUT4, GLUT1, and INSRβ protein levels in 2D myotubes differentiated in high-glucose DMEM (HGD) and MEM, with and without 1.72 µM insulin (Ins; **Figure 1A**). Relative protein abundance was assessed by normalizing values to the HGD + Ins condition, the standard in the field. HGD, a nutrient-rich medium, contains supraphysiological levels of amino acid and glucose, whereas MEM provides a composition more closely approximating physiological conditions (Cantor, 2019). In the absence of added insulin, differentiation media contains only trace amounts arising from the horse serum, and representing physiological conditions, while the addition of 1.72 µM insulin creates a supraphysiological environment. Under these differentiation conditions, insulin significantly increased GLUT4 abundance (HGD + Ins: 1.00 ± 0.00 vs. HGD: 0.41 ± 0.19, *p* = 0.021; MEM + Ins: 0.15 ± 0.13 vs. MEM: 0.74 ± 0.30, *p* = 0.021), but nutrient composition had no effect **(Figure 1B)**. In contrast, GLUT1 abundance remained unchanged across insulin and nutrient conditions **(Figure 1C)**. As a result, the GLUT4:GLUT1 ratio was ∼7.8-fold-higher in 2D myotubes differentiated in MEM with insulin compared to MEM alone (MEM + Ins: 1.32 ± 0.59 vs. MEM: 0.17 ± 0.16*, p* = 0.009; **Figure 1D**). INSRß abundance trended downward with insulin supplementation during differentiation, although the change was not statistically significant (HGD + Ins: 1.00 ± 0.00 vs. HGD: 2.09 ± 0.64, *p* = 0.114; MEM + Ins: 0.95 ± 0.12 vs. MEM: 2.01 ± 0.78, *p* = 0.124; **Figure 1E**). Together, these findings indicate that insulin supplementation during differentiation modulates glucose transporter content, with supraphysiological insulin exposure increasing GLUT4 abundance and elevating the GLUT4:GLUT1 ratio in 2D myotubes.

**Figure 1.**
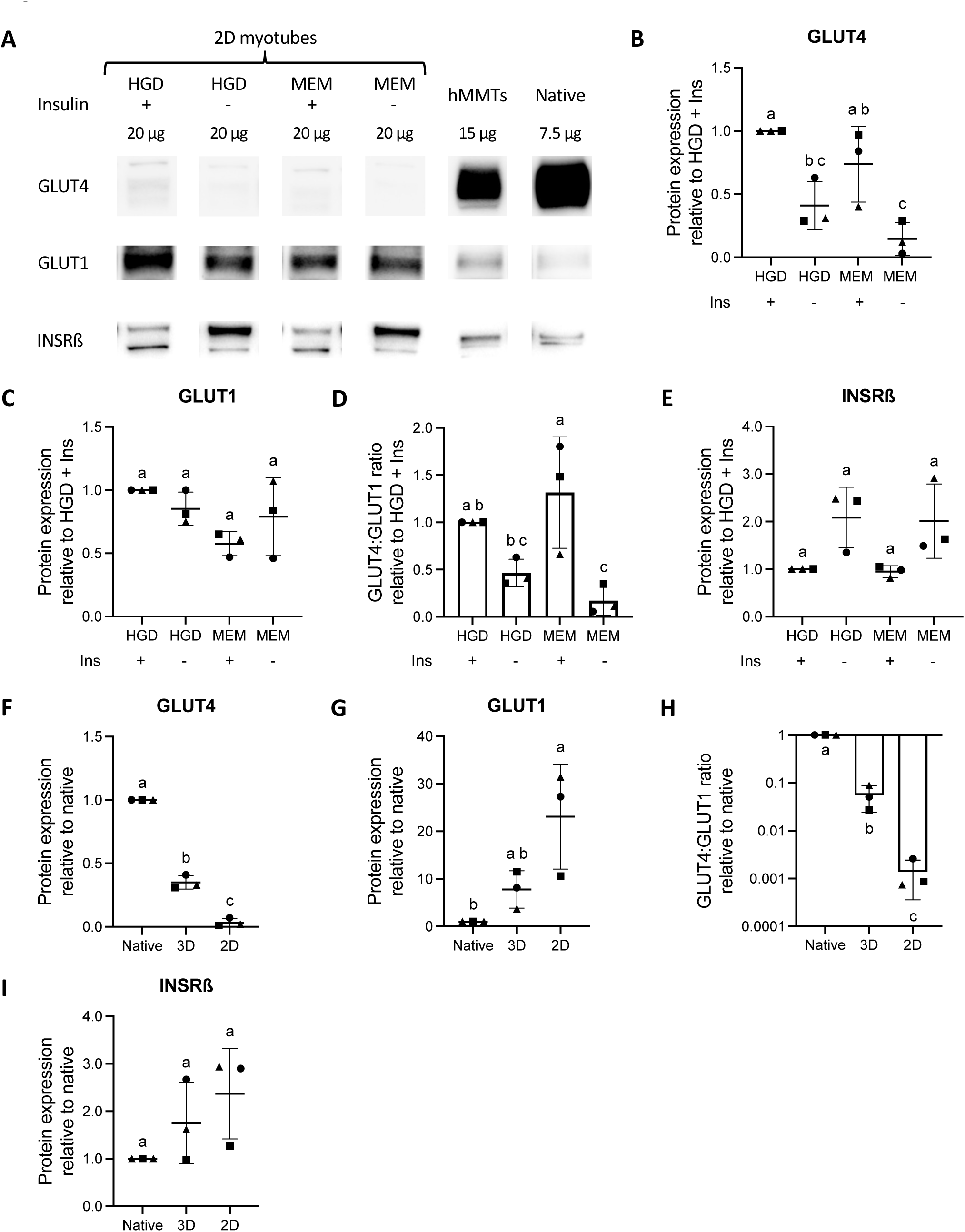
Comparison of glucose transporter and insulin receptor abundance in 2D myotubes, human skeletal muscle microtissues (hMMTs), and native human skeletal muscle. **(A)** Representative immunoblots of glucose transporter type 4 (GLUT4), glucose transporter type 1 (GLUT1), and insulin receptor ß (INSRß) in 2D myotubes, hMMTs, and native human skeletal muscle (native). Myotubes were differentiated in high-glucose Dulbecco’s Modified Eagle’s Medium (HGD; nutrient-rich) and Minimum Essential Medium (MEM; basal), with or without 1.72 µM added insulin (ins). hMMTs were differentiated in HGD with 1.72 µM insulin. Quantification of the GLUT4 **(B)** and GLUT1 **(C)** abundance, the GLUT4:GLUT1 ratio **(D)**, and INSRß **(E)** abundance in 2D myotubes cultured under the four differentiation conditions. Quantification of GLUT4 **(F)** and GLUT1 **(G)** abundance, the GLUT4:GLUT1 ratio (log_10_ scale; **H**), and INSRß **(I)** abundance in myotubes (2D), hMMTs (3D), and native human skeletal muscle; both myotubes and hMMTs differentiated in HGD with 1.72 µM insulin. n = 1-2 wells of myotubes and n = 3-6 hMMTs per condition across N = 3 independent biological replicates. The same native human skeletal muscle sample was used in all experiments. Graphs display means ± SD, with pairwise comparisons visualized using compact letter display. Circular, triangular, and square markers represent values from the first, second and third biological replicates, respectively. Statistical analysis was performed using one-way ANOVA followed by Tukey’s test.

Next, to determine how glucose transporter and insulin receptor content compares in 2D and 3D cultures, we quantified GLUT4, GLUT1, and INSRβ protein levels in myotubes (2D) and hMMTs (3D) differentiated in HGD with 1.72 µM insulin, relative to native human skeletal muscle (native; **Figure 1A**). GLUT4 abundance was ∼12-fold-higher in hMMTs than in 2D myotubes (3D: 0.35 ± 0.05 vs. 2D: 0.03 ± 0.03, *p* < 0.0001), but both remained lower than in native muscle (*p* < 0.0001; **Figure 1F**). GLUT1 levels trended lower in hMMTs than in 2D myotubes, although this difference was not statistically significant (3D: 7.78 ± 3.92 vs. 2D: 23.10 ± 11.04, *p* = 0.072; **Figure 1G**). Compared to native muscle, GLUT1 levels were higher in 2D myotubes (*p* = 0.017), whereas levels in hMMTs were not statistically different (*p* = 0.481). The GLUT4:GLUT1 ratio was ∼60-fold higher in hMMTs compared to 2D myotubes (3D: 0.06 ± 0.03 vs. 2D: 0.001 ± 0.00, *p* = 0.023), yet both remained lower than that of native muscle (*p* < 0.0001; **Figure 1H**). INSRß abundance did not differ statistically among 2D cultures, hMMTs, and native muscle **(Figure 1I)**. Thus, while GLUT4 abundance and the GLUT4:GLUT1 ratio in hMMTs remains lower than in native human skeletal muscle, both are significantly elevated compared to 2D myotubes.

### 3.2 Differentiation with added insulin enhances nuclear accretion and contractile function

To evaluate the impact of nutrient and insulin conditions on myotube fusion, we quantified the nuclear fusion index and SAA surface coverage of 2D myotubes differentiated in HGD and MEM, with or without 1.72 µM insulin. Immunostaining for SAA and Hoechst counterstaining revealed that insulin supplementation promoted nuclear accretion in 2D myotubes **(Figure 2A)**. The fusion index was significantly higher in insulin-supplemented conditions (HGD + Ins: 52.98 ± 4.53 vs. HGD: 39.01 ± 2.98, *p* = 0.004; MEM + Ins: 47.40 ± 10.00 vs. MEM: 32.78 ± 5.05, *p* = 0.002; **Figure 2B**), as was SAA surface coverage (HGD + Ins: 38.82 ± 3.32 % vs. HGD: 30.18 ± 6.13 %, *p* = 0.015; MEM + Ins: 38.42 ± 4.44 % vs. MEM: 25.37 ± 4.61 %, *p* = 0.0003; **Figure 2C**). However, neither metric was affected by the nutrient composition of the differentiation media. These findings indicate that supraphysiological insulin concentrations enhance nuclear fusion in 2D cultures, whereas the increased nutrient abundance in HGD does not.

**Figure 2.**
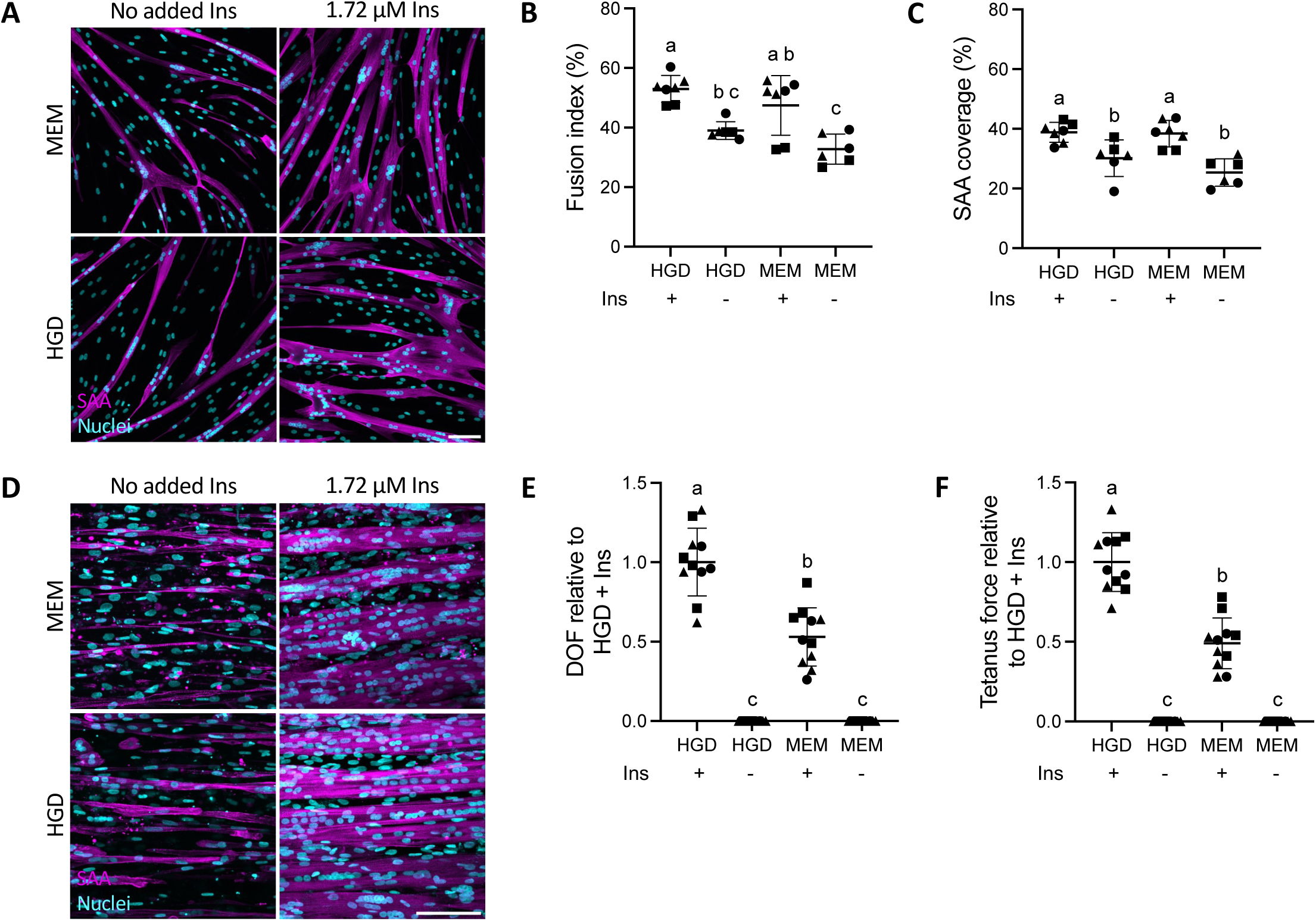
Differentiation of 2D myotubes and 3D human skeletal muscle microtissues (hMMTs) in basal vs nutrient-rich media ± insulin. **(A)** Representative 20x confocal images of 2D myotubes differentiated in high-glucose Dulbecco’s Modified Eagle’s Medium (HGD; nutrient-rich) and Minimum Essential Medium (MEM; basal), with or without 1.72 µM added insulin (ins). Sarcomeric α-actinin (SAA) is shown in magenta, and Hoechst 3342 in cyan. Scale bar = 100 µm. Dot plot quantification of myotube nuclear fusion index **(B)** and SAA surface coverage **(C)** for individual wells across differentiation conditions. n = 2-3 wells per condition across N = 3 independent biological replicates. **(D)** Representative 40x confocal images of hMMT myotubes differentiated in HGD and MEM, with or without 1.72 µM added insulin. SAA is shown in magenta, and Hoechst 3342 in cyan. Scale bar = 100 µm. Dot plot quantification of normalized dynamic oscillation of force (DOF; **E**) and tetanus force **(F)** generated by hMMTs across differentiation conditions. n = 3-4 hMMTs per condition across N = 3 independent biological replicates. Graphs display means ± SD, with pairwise comparisons visualized using compact letter display. Circular, triangular, and square markers represent technical replicates from the first, second, and third biological replicates, respectively. Data were analyzed using one-way ANOVA followed by Tukey’s test.

Next, the influence of nutrient and insulin culture conditions on contractile function was assessed by measuring the contractile force of hMMTs differentiated in HGD and MEM, with or without 1.72 µM insulin. To ensure that contractile force measurements were not affected by suboptimal cell seeding density, the optimal number of cells per tissue was first determined. While previous studies utilized a seeding density of 225K primary myoblasts per tissue (Afshar et al., 2020), reducing this number to 175 K did not compromise microtissue strength **(Supplemental Figure 1)**. Therefore, all hMMTs in this study were seeded with 175K cells per tissue. Following differentiation, immunostaining for SAA and Hoechst counterstaining confirmed the formation of multi-nucleated, striated, and aligned myotubes in insulin-supplemented conditions **(Figure 2D)**. In contrast, differentiation in the absence of added insulin resulted in sparse myotube formation, with frequent fragmentation. Consequently, hMMTs differentiated without insulin in both HGD and MEM failed to produce detectable post movement in response to EFS **(Figure 2E,F)**. Among insulin-supplemented conditions, hMMTs differentiated in HGD exhibited significantly greater DOF compared to those in MEM, with force values normalized relative to the HGD + Ins condition (HGD + Ins: 1.00 ± 0.21 vs. MEM + Ins: 0.53 ± 0.18, *p* < 0.001; **Figure 2E**). A similar trend was observed for tetanus force, which was significantly higher for differentiation in HGD compared to MEM (HGD + Ins: 1.00 ± 0.18 vs. MEM + Ins: 0.49 ± 0.16, p < 0.0001; **Figure 2F**). Thus, insulin supplementation is essential for the development of contractile hMMTs, with differentiation in nutrient-rich conditions further enhancing contractile strength.

### 3.3 hMMT peak force production requires differentiation in supraphysiological levels of insulin

Optimizing hMMT culture conditions to better mimic the physiological metabolic environment enhances their utility for modeling *in vivo* disease states. Thus, we next aimed to determine if insulin and glucose levels during differentiation could align more closely with in vivo conditions without impairing hMMT contractile function. First, the impact of reducing insulin supplementation was assessed by differentiating hMMTs in HGD with 0 µM, 0.02 µM, 0.86 µM, 1.29 µM, and 1.72 µM insulin **(Figure 3A)**. Lowering insulin levels by half had no effect on hMMT DOF **(Figure 3B)** or tetanus force **(Figure 3C)**, with values normalized to the 1.72 µM insulin condition. However, further reduction to 0.02 µM insulin— which remains supraphysiological— significantly decreased both DOF (1.72 µM: 1.00 ± 0.20 vs. 0.02 µM: 0.53 ± 0.24, *p* = 0.0004) and tetanus force (1.72 µM: 1.00 ± 0.17 vs. 0.02 µM: 0.53 ± 0.18, *p* < 0.0001). Next, the effect of reducing glucose availability was assessed by differentiating hMMTs with 0.86 µM insulin in HGD and low-glucose DMEM (LGD), with LGD containing physiological glucose levels. Immunostaining for SAA and Hoechst counterstaining confirmed the formation of multi-nucleated, striated, and aligned myotubes in both conditions **(Figure 3D)**. Despite comparable myotube morphology, differentiation in LGD yielded stronger hMMTs, with higher DOF (HGD: 1.00 ± 0.18 vs. LGD: 1.58 ± 0.66, *p* = 0.0003; **Figure 3E**) and tetanus force (HGD: 1.00 ± 0.19 vs. LGD: 1.48 ± 0.39, *p* = 0.029; **Figure 3F**), with values normalized to the HGD condition. These findings indicate that hMMTs can be differentiated in physiological glucose levels without losing contractile strength but insulin at supraphysiological concentrations is required.

**Figure 3.**
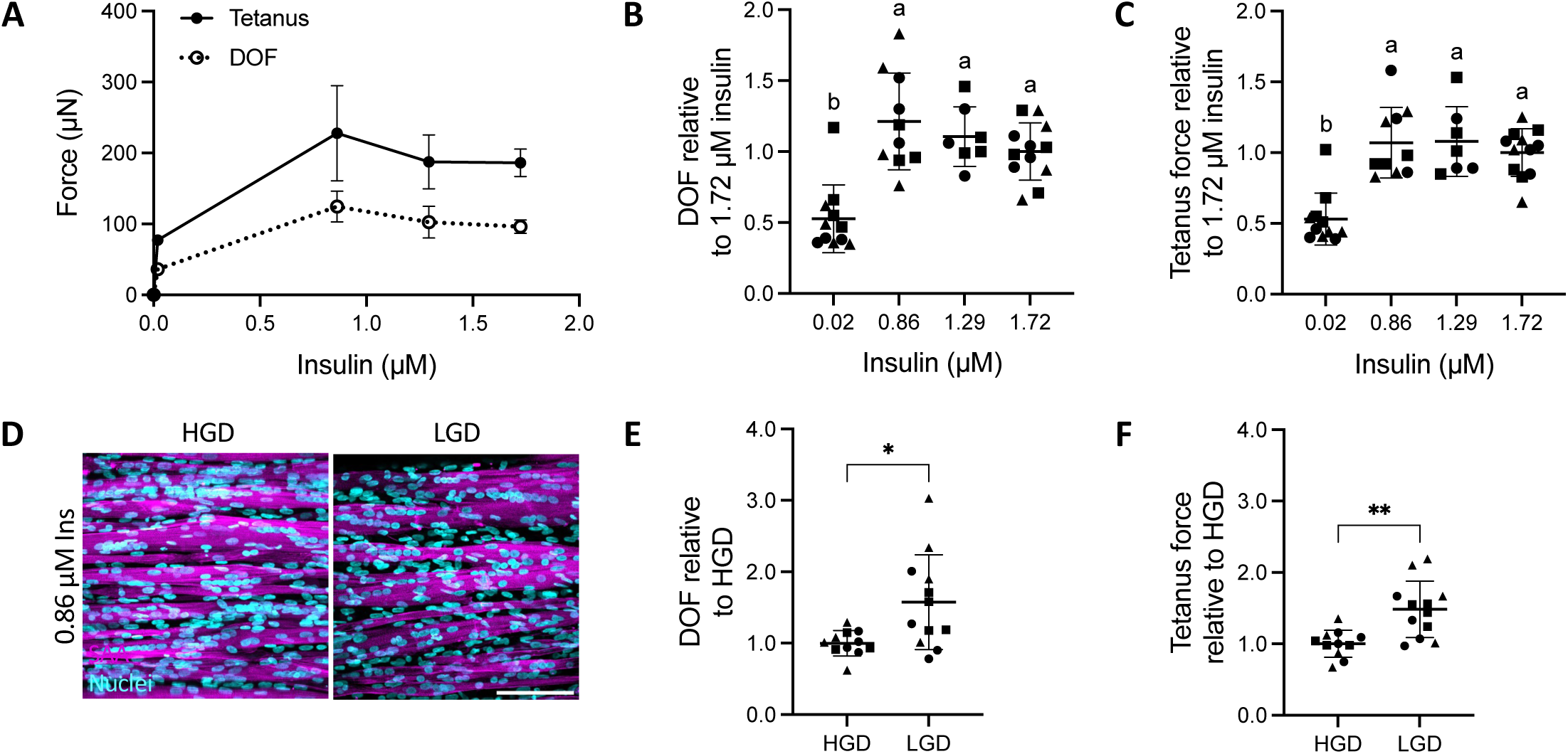
Contractile function of human skeletal muscle microtissues (hMMTs) differentiated in physiologically relevant levels of insulin and glucose. **(A)** Representative line graph depicting the dynamic oscillation of force (DOF) and tetanus force generated by hMMTs differentiated in high-glucose Dulbecco’s Modified Eagle’s Medium (DMEM) supplemented with 0 µM, 0.02 µM, 0.86 µM, 1.29 µM, and 1.72 µM insulin from a single biological replicate. Dot plots quantifying normalized DOF **(B)** and tetanus force **(C)** generated by hMMTs differentiated at each insulin concentration for all biological replicates. n = 3-4 hMMTs per insulin concentration across N = 3 independent biological replicates, except for differentiation with 1.29 µM insulin, where N = 2. Statistical analysis was performed using one-way ANOVA followed by Tukey’s test, with pairwise comparisons visualized using compact letter display. **(D)** Representative 40x confocal images of hMMT myotubes differentiated in high-glucose DMEM (HGD) and low-glucose DMEM (LGD) with 0.86 µM insulin. Sarcomeric α-actinin (SAA) is shown in magenta, and Hoechst 3342 in cyan. Scale bar = 100 µm. Dot plots quantifying normalized DOF **(E)** and tetanus force **(F)** generated by hMMTs differentiated in LGD and HGD with 0.86 µM insulin. n = 3-4 hMMTs per conditions across N = 3 biological replicates. Statistical analysis was performed using an unpaired, two-tailed *t*-test; * p ≤ 0.05, ** p ≤ 0.01. Graphs display means ± SD. Circular, triangular, and square markers represent technical replicates from the first, second, and third biological replicates, respectively.

### 3.4 Insulin-stimulated glucose uptake is enhanced in hMMTs following insulin re-sensitization

Modelling T2D in hMMTs requires the ability to capture both-insulin resistant and insulin-sensitive states *in vitro*. Differentiation under supraphysiological insulin conditions renders hMMTs largely unresponsive to insulin, necessitating its resensitization (RS). To address this, hMMTs were cultured in LGD without insulin supplementation for a four day period following the first 10 days of differentiation in LGD with 0.86 µM insulin **(Figure 4A)**. As an insulin-resistant control (CTRL), hMMTs were maintained in HGD with 1.72 µM insulin for the same duration. Immunostaining for SAA and Hoechst counterstaining revealed myotubes within RS hMMTs which appeared atrophied, with smaller myotube diameters **(Figure 4B)**. This morphological change corresponded with reduced contractile function, as RS hMMTs generated less DOF (CTRL: 1.00 ± 0.36 vs. RS: 0.40 ± 0.20, *p* < 0.0001; **Figure 4C**) and tetanus force (CTRL: 1:00 ± 0.25 vs. RS: 0.50 ± 0.18, *p* < 0.0001; **Figure 4D**), normalized to CTRL values. RS hMMTs also showed reduced fatigue resistance (CTRL: 64.68 ± 8.76 % vs. RS: 51.02 ± 14.88, *p* = 0.015; **Figure 4E**). However, despite this decline in contractility, insulin-stimulated 2DG uptake increased ∼1.08 ± 0.17-fold in CTRL hMMTs relative to basal levels, whereas RS hMMTs exhibited a ∼1.51 ± 0.20-fold increase **(Figure 4F)**. Thus, insulin withdrawal for four days post-differentiation enhances the insulin responsiveness of hMMTs but compromises their contractile function.

**Figure 4.**
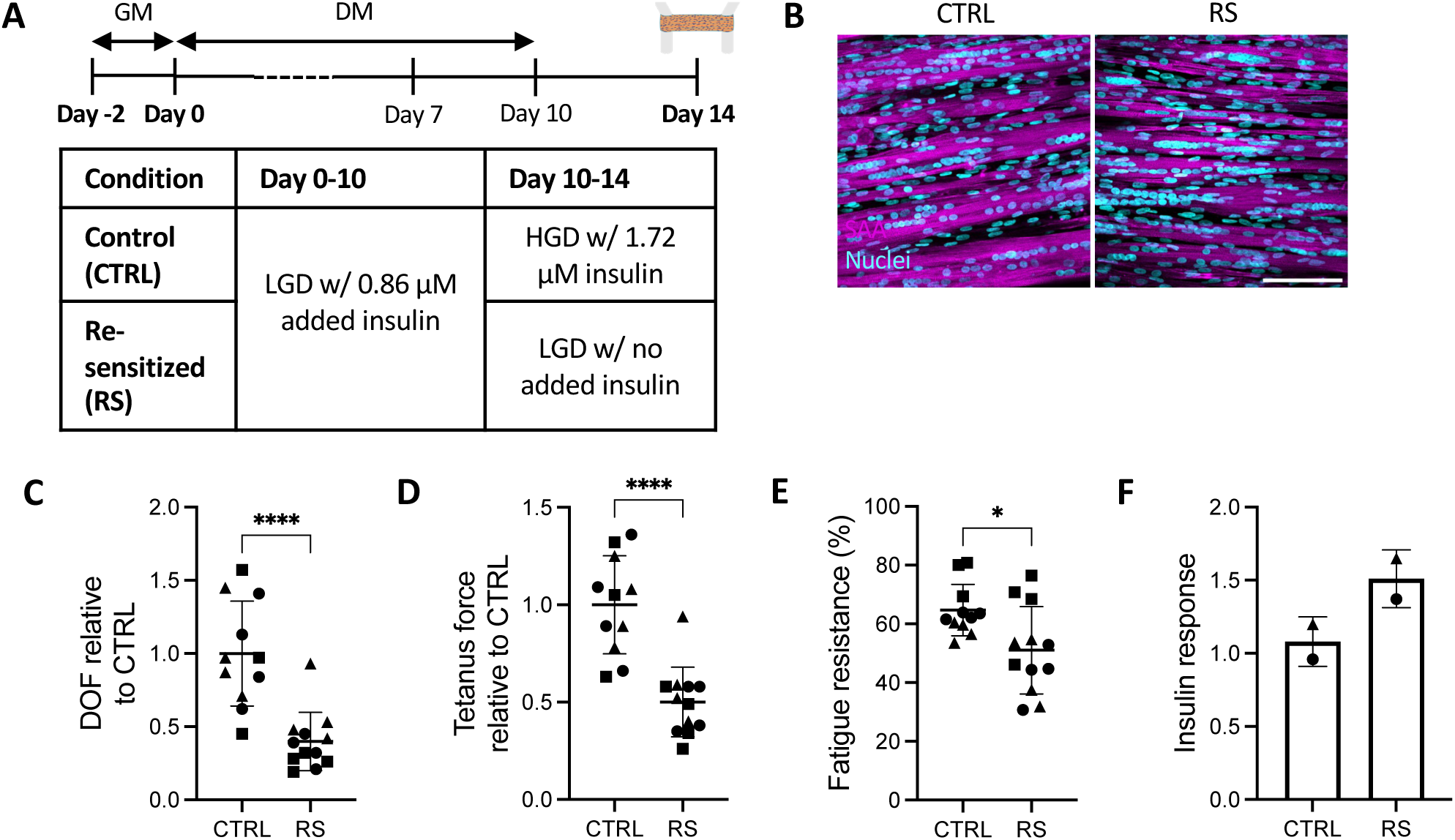
Contractile function and insulin sensitivity of human skeletal muscle microtissues (hMMTs) post insulin re-sensitization. **(A)** Schematic overview of the culture timeline for insulin re-sensitized hMMTs. hMMTs were cultured in after seeding growth media (GM) for 2 days, followed by 10 days in differentiation media (DM). For the final 4 days, they were maintained in either control (CTRL) or insulin re-sensitization (RS) media conditions. **(B)** Representative 40x confocal images of myotubes formed in CTRL and RS hMMTs. Sarcomeric α-actinin (SAA) is shown in magenta, and Hoechst 3342 in cyan. Scale bar = 100 µm. Dot plot quantification of normalized dynamic oscillation of force (DOF; **C**) and tetanus force **(D)** generated by CTRL and RS hMMTs, as well as their fatigue resistance during tetanus stimulation **(E)**. n = 3-4 hMMTs per condition across N = 3 independent biological replicates. Data were analyzed using unpaired, two-tailed T-test; * p ≤ 0.05, **** p ≤ 0.0001. **(F)** Fold change in insulin-stimulated 2DG uptake by CTRL and RS hMMTs. n = 4 hMMTs per conditions across N = 2 biological replicates. Graphs display means ± SD. Circular, triangular, and square markers represent technical replicates from the first, second, and third biological replicates, respectively.

## 4. Discussion

In this study, we investigated the suitability of human engineered skeletal muscle to model T2D-related metabolic dysfunction. While GLUT1 levels in hMMTs trended downward, albeit not significantly **(Figure 1G)**, hMMTs exhibited a ∼12-fold greater GLUT4 abundance **(Figure 1F)** and a ∼60-fold higher GLUT4:GLUT1 ratio compared to 2D myotubes **(Figure 1H)**. Although GLUT4 levels in hMMTs remained lower than those in native muscle, these findings highlight the importance of incorporating appropriate mechanical and structural cues *in vitro* to better approximate the metabolic properties of skeletal muscle. Nonetheless, we also identify a challenge in generating a physiologically relevant model of T2D from hMMTs differentiated with a standard insulin-based protocol. Differentiation without insulin supplementation resulted in hMMTs with sparse myotubes incapable of producing detectable post deflection during contraction **(Figure 2D-F)**, whereas supraphysiological insulin levels were required to generate maximally contractile hMMTs **(Figure 3A-C)**. Additionally, while hMMTs retained contractile function when cultured in physiological glucose conditions **(Figure 3E,F)**, supraphysiological amino acid levels were required **(Figure 2E,F)**, likely due to their role in promoting myotube hypertrophy. Insulin withdrawal after 10 days of differentiation restored insulin responsiveness, with ∼1.5-fold increase in insulin-stimulated glucose uptake relative to basal levels **(Figure 4F)**. However, this improvement came at the cost of reduced contractile strength and fatigue resistance **(Figure 4C-E)**, suggesting that insulin was not only necessary for optimal differentiation, but also for preserving contractile function post-differentiation. Collectively, these findings underscore the need for a robust, insulin-free differentiation protocol to facilitate the study of both cell-autonomous and insulin-induced skeletal muscle metabolic dysfunction in engineered muscle models and develop effective therapeutic strategies.

We found that hMMTs contained ∼1/3^rd^ of the GLUT4 content present in native skeletal muscle, whereas 2D myotubes derived from the same primary cell line contained only ∼1/30^th^, a finding that could be reinforced in future studies by evaluating multiple donor cell lines. This greater GLUT4 abundance in hMMTs compared to 2D myotubes likely accounts for the higher insulin responsiveness previously reported for engineered muscle (Kondash et al., 2020). Although lower GLUT4 levels in 2D myotubes reduce insulin-stimulated glucose transport, the fundamental mechanisms governing glucose uptake remain intact (Al-Khalili et al., 2003). However, this does not preclude the possibility that the conditions under which insulin resistance develops, or the mechanisms driving it, differ between 2D cultures and native muscle. Interestingly, insulin supplementation in 2D myotubes led to increased nuclear accretion **(Figure 2B,C)**, which correlated with elevated GLUT4 levels **(Figure 1B)**. This suggests that GLUT4 expression progressively increases as myoblasts differentiate, aligning with findings in trout myoblasts, where *GLUT4* mRNA expression increased throughout differentiation and was upregulated after just six hours of insulin exposure (Díaz et al., 2009).

Our findings demonstrated that lowering insulin supplementation from the standard 1.72 µM to 20 nM during differentiation significantly compromised hMMT contractile function. Consequently, we differentiated hMMTs with half the standard insulin concentration, which preserved force production. In contrast, Kondash *et al*. (2020) reported that reducing insulin supplementation from the standard 1.72 µM to 10 nM during differentiation of engineered skeletal muscle had no effect. Although 10 nM insulin is closer to physiological serum levels, it remains supraphysiological compared to concentrations observed in healthy individuals (fasting ∼0.06 nM, peaking at ∼0.42 nM postprandially) and in obese, hyperinsulinemic subjects (fasting ∼0.14 nM, peaking at ∼0.84 nM) (Polonsky et al., 1988). Despite these differences, insulin withdrawal in both our study and that reported by Kondash *et al*. resulted in a ∼1.5-fold increase in insulin-stimulated glucose uptake. While this increase is modest compared to the ∼10-fold response reported for human skeletal muscle *in vivo* (DeFronzo et al., 1985), it aligns more closely with the ∼2.0–2.5-fold response observed for isolated native muscle (Dohm et al., 1988; Lund et al., 1997). This difference suggests that the absence of innervation in engineered models may be a major contributor to the attenuated insulin response.

A model of hyperinsulinemia in engineered skeletal muscle would ideally entail a diseased and control condition that differ only in hyperinsulinemia-induced T2D-related metabolic dysfunction. However, the impairment of hMMT contractility following insulin withdrawal suggests that additional changes beyond metabolic dysfunction would occur in such a model, introducing confounding variables. Capturing cell-autonomous diabetic phenotypes using hMMTs derived from patient primary or iPSC-derived myoblasts may also be challenging, as differentiation under supraphysiological insulin levels could obscure native gene expression patterns and epigenetic signatures. These concerns can be mitigated by establishing an insulin-free differentiation protocol. Interestingly, insulin-like growth factor I (IGF-I) and insulin function as potent fetal skeletal muscle growth factors (Brown, 2014). However, fetal muscle formation is regulated through an autocrine/paracrine system involving insulin growth factors and their binding proteins (Oksbjerg et al., 2004). IGF-I promotes commitment of cells to the muscle lineage, and cells secrete increasing levels of insulin-like growth factor II (IGF-II) during muscle differentiation, which in turn promotes fusion into multinucleated myofibers (Aboalola & Han, 2017). Through a dual regulatory mechanism, IGF-II suppresses IGF-I gene expression, and upregulates its own expression (Jiao et al., 2013). Primary human myoblasts differentiated in serum- and insulin-free medium supplemented with IGF-I formed thicker multinucleated myotubes in 2D cultures that were responsive to electrical pulse stimulation, exhibited increased GLUT4 expression, and showed enhanced insulin-stimulated glucose uptake compared to cultures differentiated without IGF-I (Dreher et al., 2024). These findings support a theory wherein an IGF-I-based differentiation protocol may offer a feasible pathway for the development of contractile hMMTs independent of insulin.

## 5. Conclusions

Our findings reveal that engineered skeletal muscle exhibits a more mature glucose transporter profile than 2D myotubes, characterized by higher GLUT4 levels and an increased GLUT4:GLUT1 ratio. The cell-cell and cell-extracellular matrix interactions present in hMMTs are likely key contributors to upregulating GLUT4 expression. To further enhance the physiological relevance of these models, electrical or optogenetic stimulation protocols (Khodabukus et al., 2019; Mills et al., 2019) can be implemented, shifting the metabolism of hMMTs closer to that of native. Currently, the reliance on standard insulin-based differentiation protocols limits the use of hMMTs for studying cell-autonomous and insulin-induced skeletal muscle metabolic dysfunction. This challenge may be overcome by leveraging the mechanisms that drive *in vivo* skeletal muscle myogenesis to generate contractile hMMTs independent of insulin. Establishing such models would not only facilitate the study of T2D-related metabolic dysfunction but also enable investigations into the role of contraction and contraction-induced myokine secretion in enhancing muscle insulin sensitivity, ultimately informing the development of novel therapeutics.

## DATA AVAILABILITY

The datasets that support the findings of this study are available from the corresponding author upon reasonable request.

## SUPPLEMENTAL MATERIAL

Supplemental Tables 1-4 and Supplemental Figure 1

## AUTHOR CONTRIBUTIONS

R.K. collected and pre-processed the cadaveric skeletal muscle tissue for this study. Y.T. and P.M.G. conceptualized the study, designed experiments, and interpreted data. Y.T. conducted research, analyzed data, and prepared figures. J.N. conducted research and analyzed data. Y.T. drafted the initial manuscript, which was then edited by P.M.G. All authors reviewed and approved the manuscript.

## Supplemental Figures

**Supplemental Figure 1.**
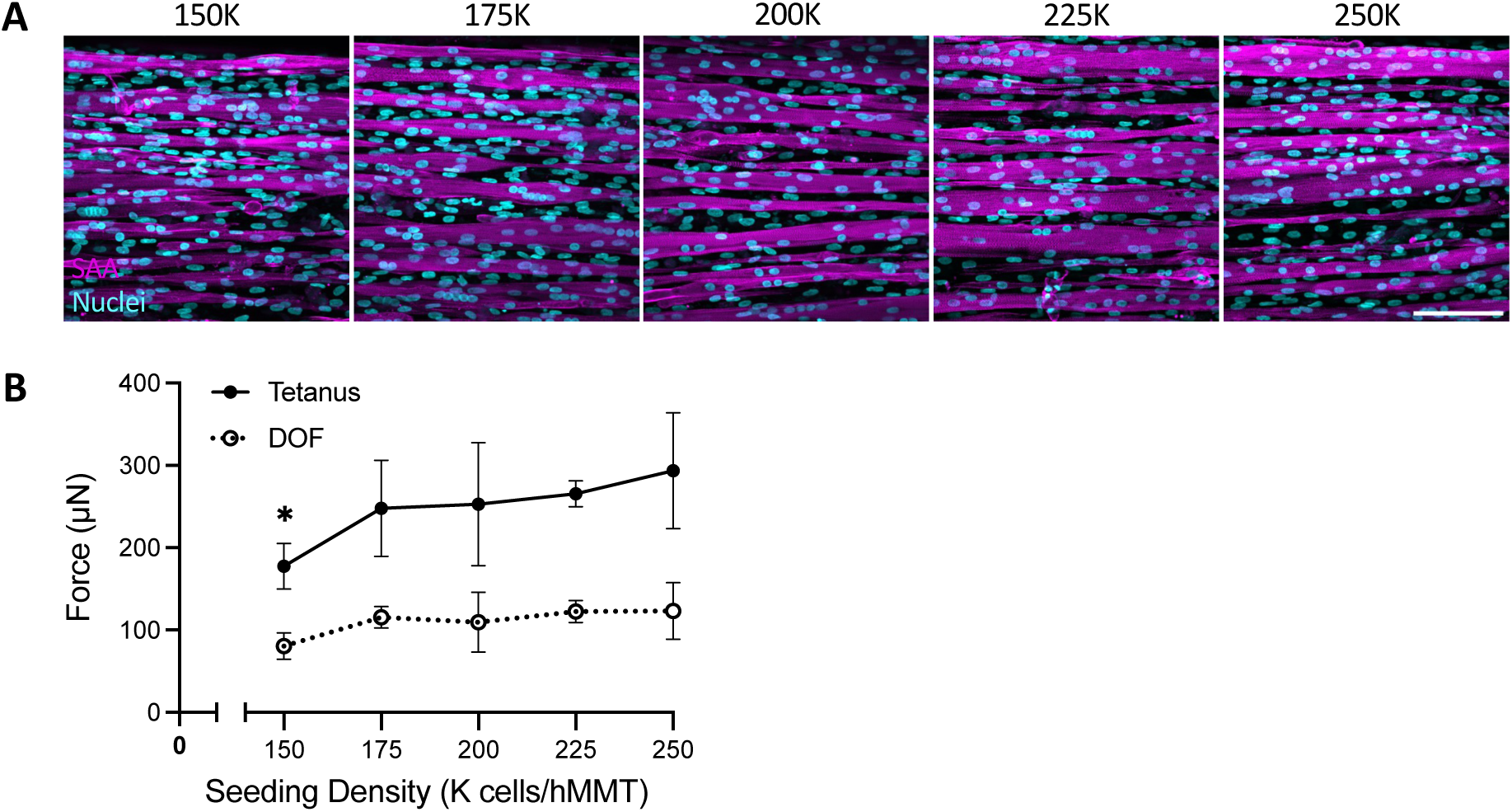
Optimization of human skeletal muscle microtissue (hMMT) seeding density to achieve peak force. **(A)** Representative 40x confocal images of myotubes formed in hMMTs seeded with 150 K, 175 K, 200 K, 225 K, and 250 K cells. hMMTs were immunostained for sarcomeric α-actinin (SAA, magenta) and counterstained with Hoechst 3342 (cyan). Scale bar = 100 µm. **(B)** Line graph of dynamic oscillation of force (DOF) and tetanus force produced by electrical field stimulation of hMMTs at each seeding density. Data show means ± SD for n = 3-4 hMMTs from N = 1 independent biological replicate. Statistical analysis was performed using one-way ANOVA followed by Dunnett’s test, with comparisons made relative to the 225 K cells/hMMT condition: * p ≤ 0.05.

**Supplemental Table 1.** Composition of media/solutions used for human primary myoblast expansion, 2D myotube differentiation, and hMMT production and culture.

| Media/solution | Composition |
| --- | --- |
| Rat tail collagen coating solution | ~16 µg/mL rat tail Collagen I in 0.1 M acetic acid solution prepared with dH<sub>2</sub>O |
| Myoblast growth medium | Ham's F-10 nutrient mix, 20 % FBS, 5 ng/mL bFGF, 1 % P/S |
| 2D differentiation medium | High glucose DMEM or MEM, 2 % HS, 1 % P/S, with or without 1.72 µM insulin |
| Fibrinogen stock solution | 10 mg/mL fibrinogen in 0.9 % (w/v) NaCl solution prepared with dH<sub>2</sub>O |
| Hydrogel master mix | High glucose DMEM (40 % v/v), 4 mg/mL fibrinogen (40 % v/v), Geltrex™ (20 % v/v) |
| After-seeding growth medium | Ham's F-10 nutrient mix, 20 % FBS, 1.5 mg/mL ACA (3 % v/v), 1 % P/S |
| 3D differentiation medium | High glucose DMEM, MEM, or low glucose DMEM, 2 % HS, 2 mg/mL ACA (4 % v/v), 1 % P/S, without or with up to 1.72 µM insulin |

**Abbreviations:** FBS = fetal bovine serum, bFGF = basic fibroblast growth factor, P/S = Penicillin-Streptomycin, DMEM = Dulbecco's Modified Eagle's Medium, MEM = Minimum Essential Medium, HS = horse serum, ACA = 6-aminocaproic acid.

**Supplemental Table 2.** Host species, working dilutions, and manufacturer information of the antibodies and counterstain used in this study.

| Antibody | Host species | Dilution | Manufacturer and catalog number |
| --- | --- | --- | --- |
| Anti-sarcomeric α-actinin | Mouse | 1:800 | Sigma-Aldrich, Cat. #A7811 |
| Alexa Fluor™ 488 anti-mouse | Goat | 1:500 | Invitrogen, Cat. #A11001 |
| Hoechst 33342 | - | 1:1000 | Invitrogen, Cat. #H3570 |
| Anti-glucose transporter type 4 | Rabbit | 1:1000 | Invitrogen, Cat #PA1-1065 |
| Anti-glucose transporter type 1 | Mouse | 1:100 | Santa Cruz Biotechnology, Cat. #sc-377228 |
| Anti-insulin receptor $\beta$ | Mouse | 1:200 | Satna Cruz Biotechnology, Cat. #sc-57342, |
| Horseradish peroxidase-conjugated anti-rabbit | Goat | 1:5000 | Cell Signaling Technology, Cat. #7074, |
| Horseradish peroxidase-conjugated anti-mouse | Horse | 1:5000 | Cell Signalling Technology, Cat. #7076 |

Abbreviations: HRP = horseradish peroxidase.

**Supplemental Table 3.** Composition of the assay solution used for measurements of 2DG uptake.

| Enzyme or stock solution | Solvent | Manufacturer and catalog number | Volume or units per 10 mL of assay solution |
| --- | --- | --- | --- |
| 200 mM TEA buffer, pH 8.1 | 200 mM KCl in dH<sub>2</sub>O | Sigma-Aldrich, Cat. #T1502 | 2.5 mL |
| 0.4 % BSA | dH<sub>2</sub>O | BioShop, Cat. #ALB005 | 500 µL |
| 10 mM NADP sodium salt | dH<sub>2</sub>O | Millipore, Cat. #481971 | 100 µL |
| Diaphorase from *C. kluyveri* | 50 mM TEA buffer, pH 8.1 | Sigma-Aldrich, Cat. #D5540 | 2 U |
| 2 mM Resazurin sodium salt | 50 mM TEA buffer, pH 8.1 | Sigma-Aldrich, Cat. #R7017 | 10 µL |
| *L. mesenteriodes* G6PDH | - | Sigma-Aldrich, Cat. #G5760 & #G8404 | 175 U |
| dH<sub>2</sub>O | - | - | Up to 10 mL |

Abbreviations: TEA = triethanolamine, BSA = bovine serum albumin, G6PDH = Glucose-6-phosphate dehydrogenase.

**Supplemental Table 4.** Number of pooled sample counts or technical replicates, and statistical tests for each experiment.

| Data presented | Figure | # of technical replicates or pooled sample counts | Statistical test |
| --- | --- | --- | --- |
| GLUT4, GLUT1, and INSRβ abundance in 2D myotubes differentiated in basal vs nutrient-rich media ± insulin | Fig. 1B-E | n = 1-2 pooled wells averaged over N = 3 experiments | One-way ANOVA + Tukey's test |
| GLUT4, GLUT1, and INSRβ abundance in 2D myotubes, hMMTs, and native skeletal muscle | Fig. 1F-I | n = 1-2 pooled wells and n = 3-6 pooled hMMTs averaged over N = 3 experiments | One-way ANOVA + Tukey's test |
| Fusion index and SAA surface coverage of 2D myotubes differentiated in basal vs nutrient-rich media ± insulin | Fig. 2B,C | n = 2-3 wells averaged over N = 3 experiments | One-way ANOVA + Tukey's test |
| DOF and tetanus force of hMMTs differentiated in basal vs nutrient-rich media ± insulin | Fig. 2E,F | n = 3-4 hMMTs averaged over N = 3 experiments | One-way ANOVA + Tukey's test |
| DOF and tetanus force of hMMTs differentiated in HGD with 0.02 μM, 0.86 μM, 1.29 μM, and 1.72 μM insulin | Fig. 3B,C | n = 3-4 hMMTs averaged over N = 3 experiments (except for 1.29 μM insulin condition where N = 2) | Unpaired, two-tailed t-test |
| DOF and tetanus force of hMMTs differentiated in HGD and LGD with 0.86 μM insulin | Fig. 3E,F | n = 3-4 hMMTs averaged over N = 3 experiments | Unpaired, two-tailed t-test |
| DOF, tetanus force, and fatigue resistance of CTRL and RS hMMTs | Fig. 4C-E | n = 3-4 hMMTs averaged over N = 3 experiments | Unpaired, two-tailed t-test |
| Insulin response of CTRL and RS hMMTs | Fig. 4F | n = 4 hMMTs averaged over N = 2 experiments | N/A |
| DOF and tetanus force of hMMTs seeded with 150 K – 250 K cells | Sup. Fig. 1 | n = 3-4 hMMTs from N = 1 experiment | One-way ANOVA + Dunnett's test |

**Abbreviation:** GLUT4 = glucose transporter type 4, GLUT1 = glucose transporter type 1, INSRβ = insulin receptor $\beta$ , hMMTs = human muscle microtissues, SAA = sarcomeric α-actinin, DOF = dynamic oscillation of force, HGD = high glucose DMEM, LGD = low glucose DMEM, CTRL = control, RS = re-sensitized.

